# Multiple horizontal acquisitions of plant genes in the whitefly *Bemisia tabaci*

**DOI:** 10.1101/2022.01.12.476015

**Authors:** Clément Gilbert, Florian Maumus

## Abstract

The extent to which horizontal gene transfer (HGT) has shaped eukaryote evolution remains an open question. Two recent studies reported four plant-like genes acquired through two HGT events by the whitefly *Bemisia tabaci*, a major agricultural pest (Lapadula et al. 2020; Xia et al. 2021). Here, we uncovered a total of 49 plant-like genes deriving from at least 24 independent HGT events in the genome of the MEAM1 whitefly. Orthologs of these genes are present in three cryptic *B. tabaci* species, they are phylogenetically nested within plant sequences, and expressed and under purifying selection. The predicted functions of these genes suggest that most of them are involved in plant-insect interactions. Thus, substantial plant-to-insect HGT may have facilitated the evolution of *B. tabaci* towards adaptation to a large host spectrum. Our study shows that eukaryote-to-eukaryote HGT may be relatively common in some lineages and it provides new candidate genes that may be targeted to improve current control strategies against whiteflies.

## Introduction

Horizontal gene transfer (HGT) is the passage of genetic material between organisms by means other than reproduction. The patterns, mechanisms and vectors of HGT are well-characterized in prokaryotes, in which these transfers are ubiquitous and a major source of innovation ^1^. In eukaryotes, HGT has long been considered anecdotal because of multiple barriers that should impede such transfers, or controversial, as resulting from phylogenetic artifacts or contaminant sequences ^2 3^. Yet, recent studies have reported robust HGT events in various eukaryotic organisms. In unicellular eukaryotes, HGT appears widespread, with dozens of foreign genes characterized in a large diversity of taxa, many of which are involved in adaptive functions ^4 2 5 6 7 8^. Increasing numbers of HGT events are also reported in multicellular eukaryotes. For example, in plants, multiple cases of prokaryote-to-plant and plant-to-plant HGT have been characterized ^9 10 11^. In metazoans, several taxa are notorious for their relatively high content in foreign genes, such as root-node nematodes ^12^ and rotifers ^13^, and HGT from both endosymbiotic and non-endosymbiotic bacteria can be common, especially in arthropods ^14 15 16 17^. While functional studies remain scarce, it appears that many bacterial and fungal genes independently acquired by nematodes and arthropod lineages are involved in adaptation to phytophagy ^18 19^. For example, a Glycosyl Hydrolase Family 32 gene acquired horizontally from rhizobial bacteria by the plant parasitic nematode *Globodera pallida* is expressed in the digestive system during feeding and is involved in metabolizing plant-derived sucrose ^20^. In the coffee borer beetle (*Hypothenemus hampei*, Coleoptera), a mannanase-encoding gene acquired from *Bacillus*-like bacteria was shown to hydrolyze the major polysaccharide of coffee berries, thus likely facilitating adaptation of the beetle to a new food source ^21^. Furthermore, in the phytophagous mite *Tetranychus urticae*, a cysteine synthase gene acquired from bacteria is involved in plant cyanide detoxification and thus likely enabled colonization of a new niche by this mite and other arthropods ^22^. Perhaps even more remarkable is the acquisition of four horizontally transferred genes (two BAHD acyltransferases called BtPMaT1 and BtPMaT2 and ribosome inactivating proteins [RIPs]) not from bacteria or fungi but directly from plants, by the sweet potato whitefly *Bemisia tabaci* ^23 24^. Whiteflies (family Aleyrodidae) are herbivorous hemipteran insects that are important agricultural pests because of their feeding habits and the many viruses they transmit to plants. Functional assays revealed that the BtPMaT1 protein detoxifies plant phenolic glucosides that are normally used by plants to protect themselves against insect herbivores. Thus, the acquisition of a plant gene by whiteflies through HT enabled them to thwart plant defenses and may in part explain why these insects have become generalist phloem-feeders ^24^. Interestingly, whiteflies feeding on transformed tomato plants expressing BtPMaT1 gene-silencing fragments showed increased mortality and reduced fecundity ^24^. Therefore, identifying genes of plant origin in herbivore insects can provide targets to engineer new pest control strategies.

## Results and discussion

### Identification of plant-to-whitefly HGT candidates in MEAM1

To assess the extent to which plant-to-insect HGT may have shaped interactions between whiteflies and their host plants, we conducted a systematic search for genes of plant origin in the *B. tabaci* MEAM1 genome ^25^. We first applied a high-throughput method to infer a last common ancestor (LCA) for each of the 15,662 predicted proteins in *B. tabaci*. To this end, we used MMseqs2 taxonomy ^26^ which proposes three different modes of LCA inference (i.e. single search LCA, approximate 2bLCA, and best hit) for a query protein by processing the taxonomy of proteins retrieved by similarity search. As target database, we used the Uniref90 from which we removed Aleyrodidae proteins in order to avoid confusion in LCA prediction (Figure 1).The current release of the Uniref90 database contains 119,222,328 protein sequences each representative of a group of UniprotKB proteins clusterized at 90% similarity threshold ^27^. The UniprotKB database itself currently contains 230,328,648 proteins from all types of organisms, including 13,238,084 proteins from plants (Viridiplantae) and 51,638,861 proteins from non-plant eukaryotes (https://www.ebi.ac.uk/uniprot/TrEMBLstats, ^28^). In our approach, the placement of a protein LCA in plants indicates potential HGT from plants to *B. tabaci* after the emergence of Aleyrodidae. The deeper the LCA, the more conserved the protein across the breadth of plant taxonomy, which reduces the risk to confuse donor and recipient taxa as LCA and the risk of LCA inference based on contamination in the target database (e.g. contaminations in genome assemblies leading to erroneous taxonomic metadata). We predicted an LCA for each of the MEAM1 proteins using MMseqs2 taxonomy with the three LCA inference protocols. As expected, the vast majority of the 15,662 MEAM1 proteins (e.g. 12,200 in the best hit LCA mode) were assigned to a metazoan LCA (Supplementary Dataset 1). Interestingly, combining the LCA predictions from the different protocols, MMseqs2 predicted plant LCA with at least one of them for 365 MEAM1 proteins. Most LCA (297/365), however, were placed at tips of the plant taxonomy (genus, species, varietas and subtribe), reflecting that the target proteins used for LCA inference are poorly conserved across plants and these LCA were considered of low confidence. By contrast, we identified 68 *B. tabaci* MEAM1 proteins for which LCA was inferred deeper than genus level in plants and which were hence considered potentially transferred from plants to Aleyrodidae. This included the two BAHD acyltransferase genes (BtPMaT1 and BtPMaT2) reported by Xia et al. <u>^24^</u> and the two RIPs identified by Lapadula et al. ^23^.

**Figure 1.**
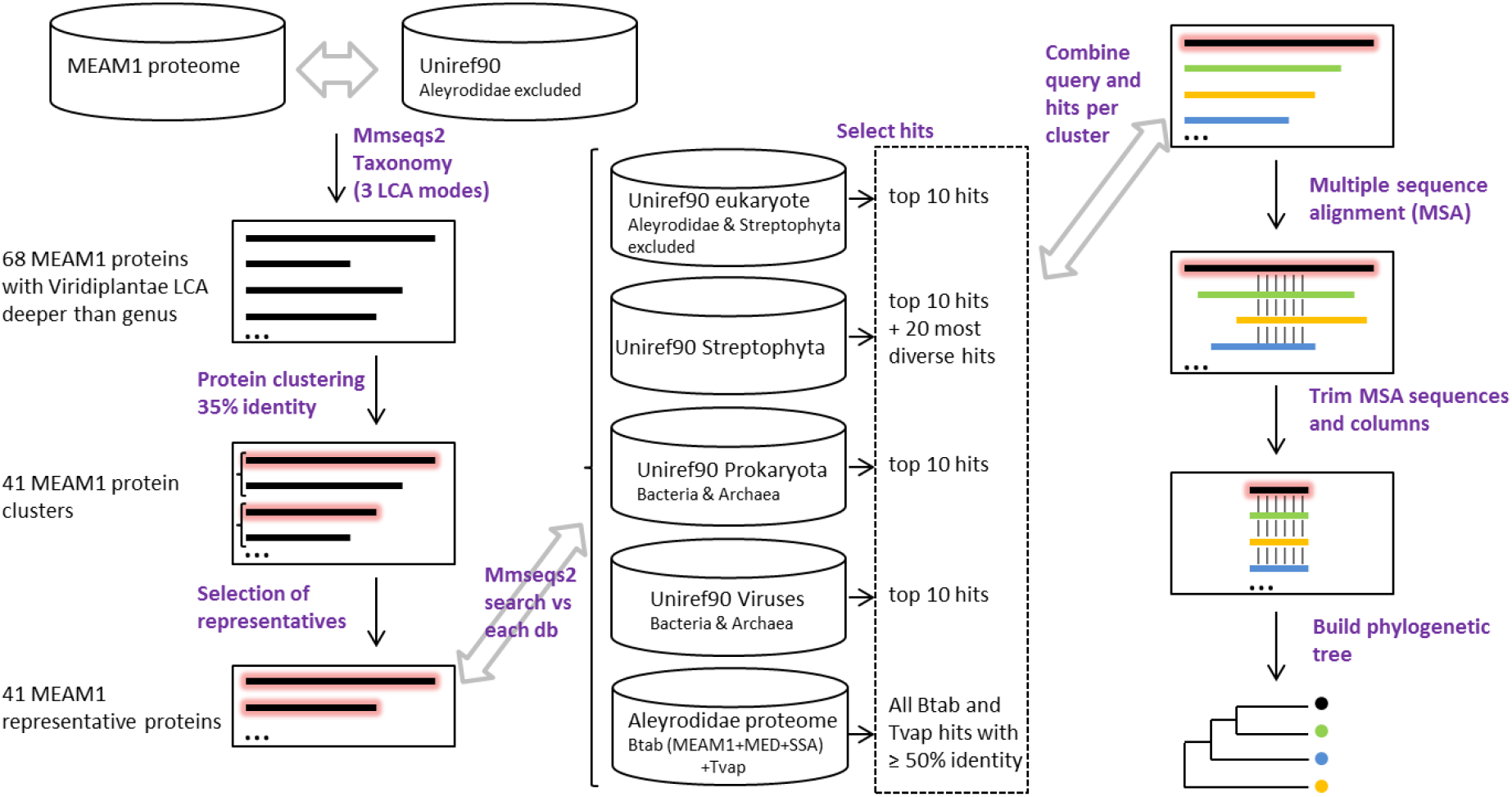
Overview of the workflow used to identify plant-derived protein candidates in the MEAM1 proteome. The containers represent target databases, the different steps are indicated in purple. The MEAM1 proteins are represented by thick black lines and the representative ones are highlighted in red.

### Assessment of plant-to-whitefly HGT candidates in MEAM1

To further characterize the taxonomic origin of candidate *B. tabaci* plant-derived genes, we grouped them in 41 clusters of orthologous proteins and used one representative sequence per cluster to perform similarity searches against all Aleyrodidae proteomes as well as against the UniRef90 protein database of streptophytes, that of eukaryotes without streptophytes and Aleyrodidae, and those of prokaryotes and viruses (see Materials and Methods). The ten best hits from each search supplemented with the twenty most diverse hits against streptophytes were used to build a multiple alignment which was submitted to phylogenetic analysis (Figure 1).The aim of this hit collection protocol was to address the relationships of each representative protein with regard to their best hits. Manual examination of the resulting trees and supporting alignments indicated that the signal observed in 17 out of 41 datasets was not in support of plant-to-Aleyrodidae gene transfer. The reason for this was either because no or too few plant hits are present in the final alignment, or the MEAM1 representative protein is significantly truncated compared to selected orthologs, or the phylogenetic placement with respect to plant orthologs is in contradiction with plant-to-whitefly HGT (see Supplementary Figure 1 and Supplementary Dataset 2). For three clusters, no tree containing the representative protein was obtained as a result of applying a filter against poorly aligned proteins. Remarkably, the topology of 24 trees was consistent with the scenario of plant-to-Aleyrodidae gene transfer. In 22 of the 24 trees, the *B. tabaci* proteins were nested within clades of proteins of streptophyte origin (Figure 2A, Supplementary Figure 2, Supplementary Dataset 3), as in ^23,24.^. In the remaining two trees (Bta13103 and Bta14885), the *B. tabaci* proteins are sister to all proteins of streptophyte origin (Figure 2B and Supplementary Figure 2). The topology of these two trees could be explained either by extensive sequence adaptation after transfer or by donor plant lineage being extinct or under-represented in genome databases. In support of a plant origin, homologous genes are absent from the proteomes of non-Aleyrodidae insects and from that of all other metazoans. Furthermore, 23 clusters are shared between MEAM1 and at least one other *B. tabaci* cryptic species (MED1 and SSA-ECA) independently sequenced in different laboratories, indicating that they are *bona fide B. tabaci* genes, and not contaminating plant genes. We further investigated the genomic environment of MEAM1 candidate plant-derived genes by assigning a taxonomic origin to each of their nearest flanking protein-coding genes. After excluding Aleyrodidae proteins from the Uniref90 database, we were able to retrieve a hit for 61 out of the 76 flanking genes investigated and found that all but two of them had a metazoan best hit (47 out of 61 having a Pancrustacea best hit) (Supplementary Table 1). These results confirm that the genes of putative plant origin stand in the context of animal contigs in the *B. tabaci* MEAM1 assembly. Altogether, these evidences indicate that as proposed for BAHD acyltransferases and RIPs ^23,24^, these 24 representative genes (Supplementary Dataset 4) were acquired by an ancestor of *B. tabaci* through plant-to-insect HGT.

**Table 1.**
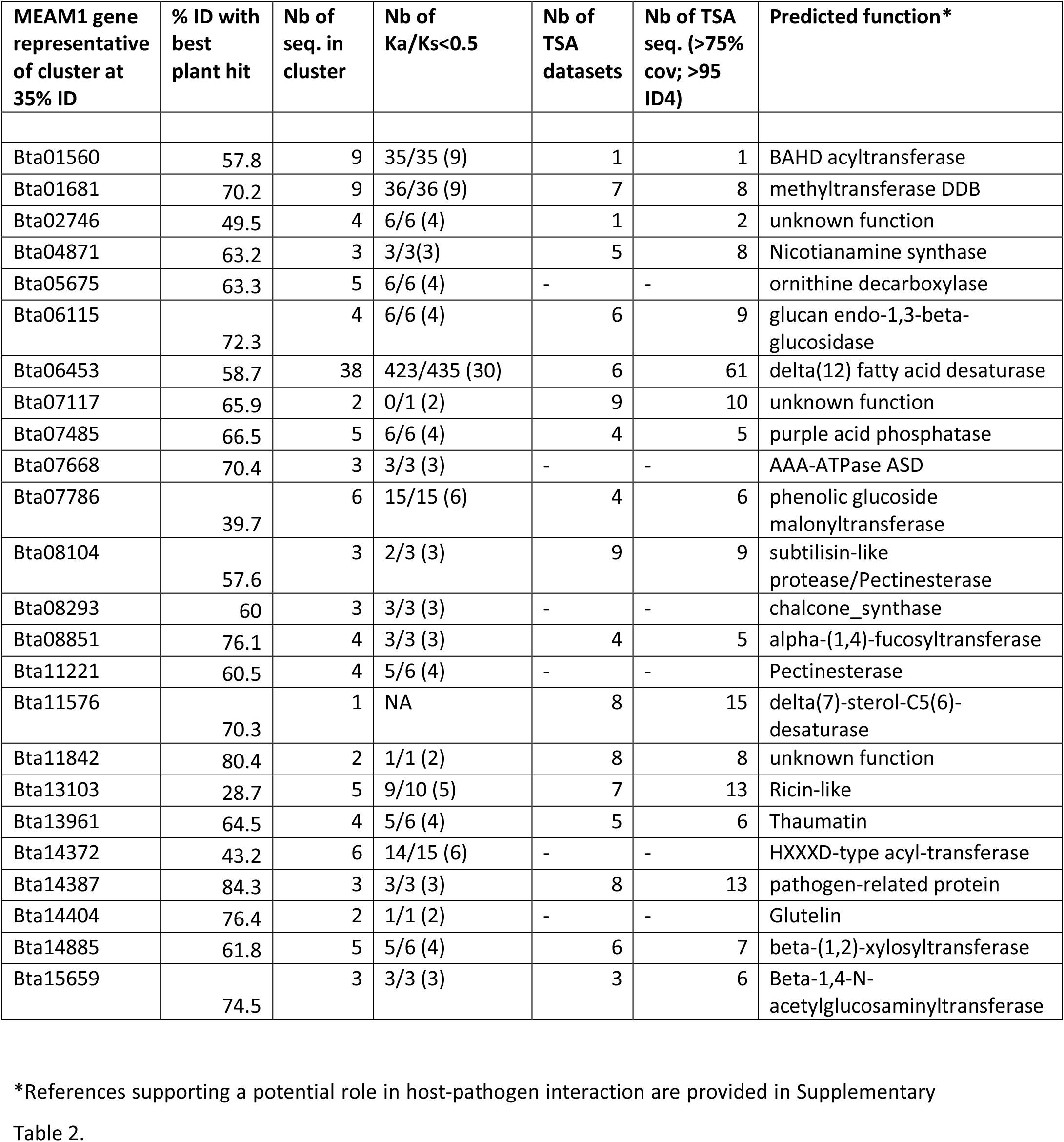
Characteristics of representative sequences of each cluster of plant-derived *Bemisia tabaci* genes.

**Figure 2.**
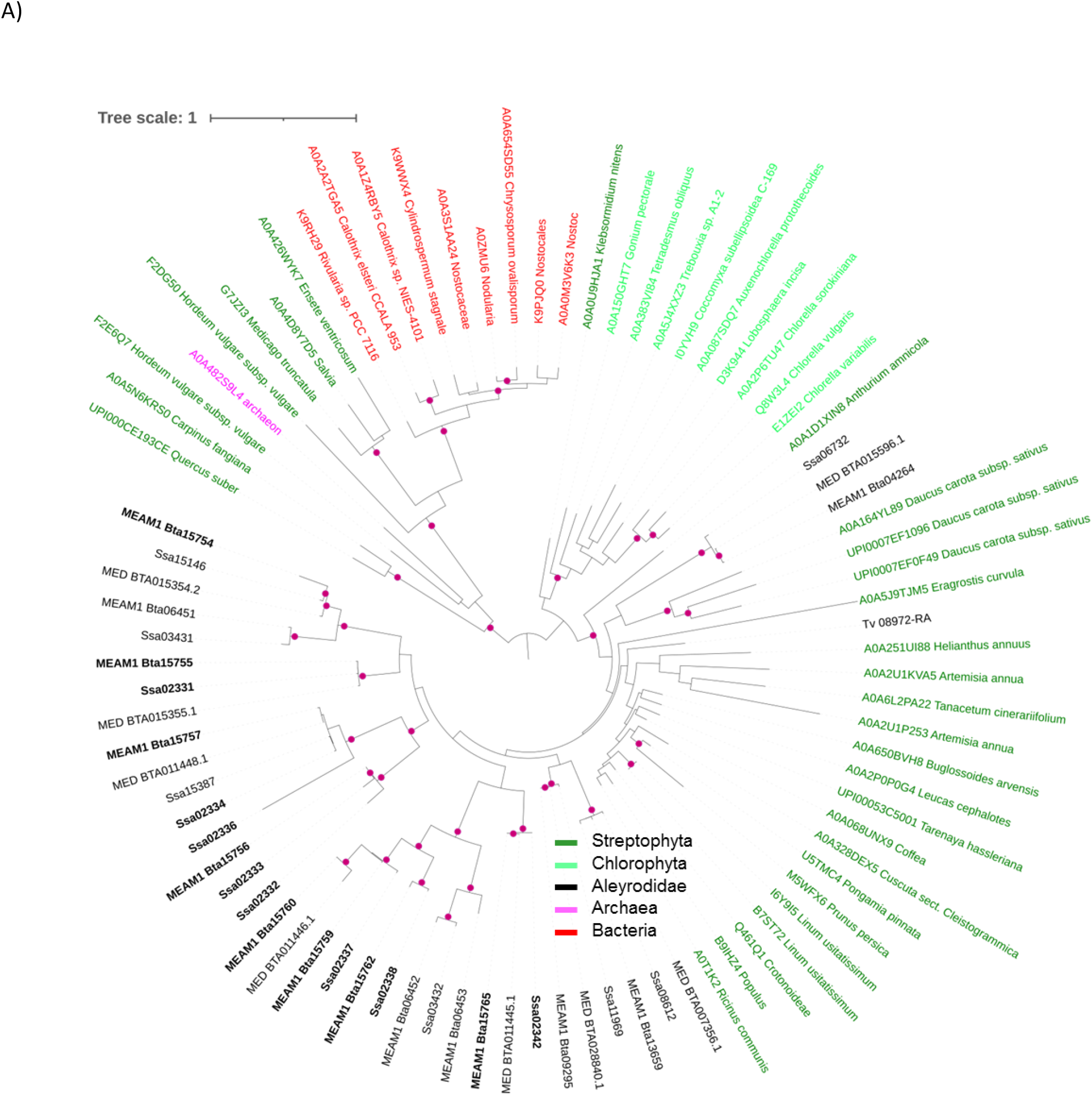
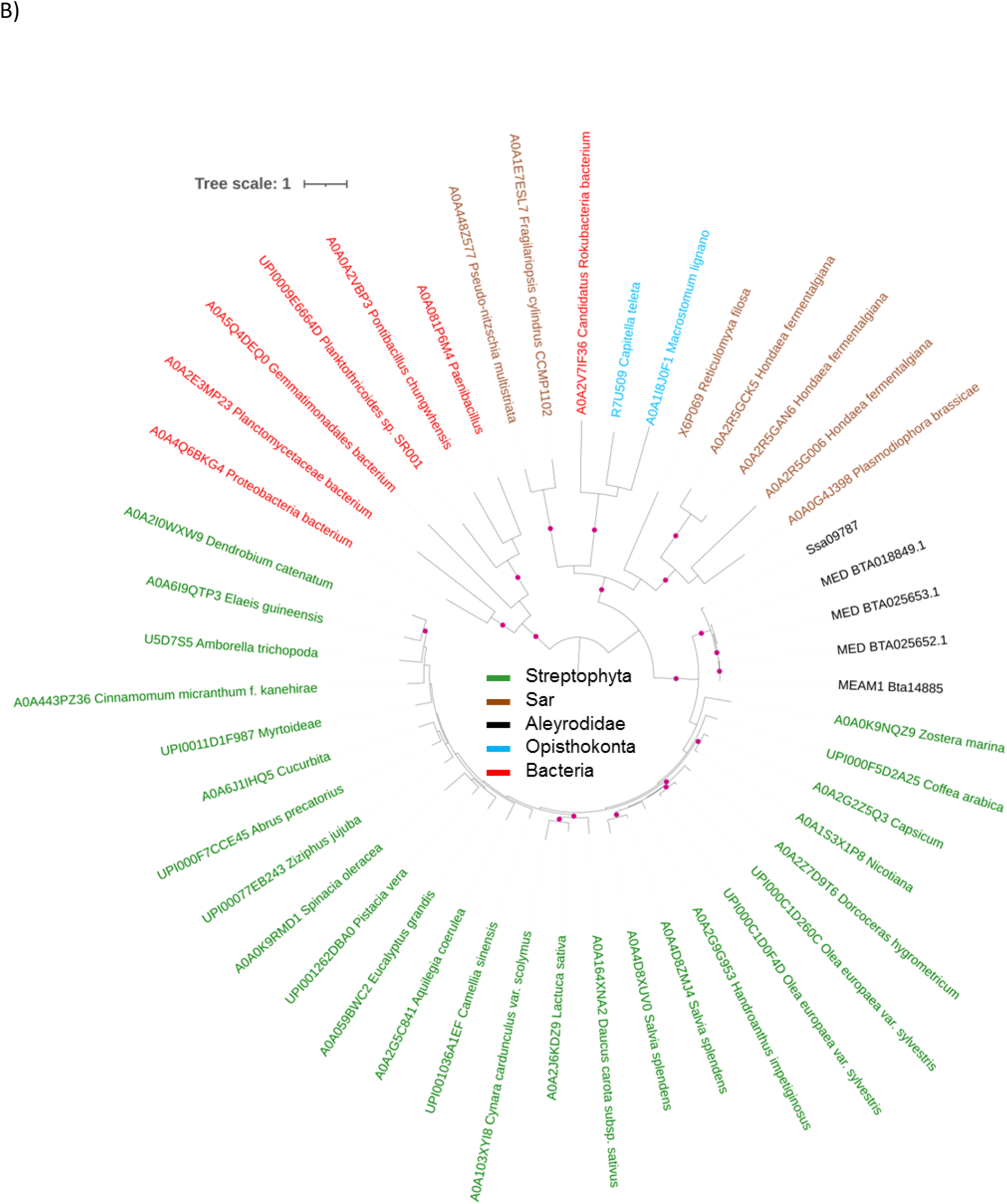
Phylogenetic relationships of potential plant-derived *Bemisia tabaci* genes. Phylogenetic trees showing the Bta06453 cluster, annotated as delta(12) fatty acid desaturase family (A) and the Bta14885 cluster, annotated as beta-(1,2)-xylosyltransferase family (B). Each tree is inferred from the multiple sequence alignment of the cluster representative sequence with its homologs found across Aleyrodidae proteins and Uniref90 proteins from eukaryotes, prokaryotes and viruses. The labels for Uniref90 proteins are either species names or the names of the first taxonomic level comprising all the proteins of a specific Uniref90 cluster. The SSA-ECA proteins are labelled as " Ssa ». In (A), the labels of the *B. tabaci* proteins present in the FAD genomic hotspots found in the MEAM1 and SSA-ECA genomes are in bold and the *Trialeurodes vaporariorum* protein is labelled as “Tv”. The bootstrap values above 70% are indicated as magenta disks. The color legend indicating the taxonomy of proteins is indicated inset. For the ease of representation, the trees are presented in a circular mode rooted at midpoint. Unrooted trees are available in the Supplementary Figures 1 & 2.

### Hypotheses alternative to horizontal transfer

A number of earlier studies reporting cases of horizontal gene transfer in animals have later been dismissed and shown to be plagued by bacterial contamination ^29^ or technical artefact in the similarity search or phylogenetic analyses ^3^. In other cases, it is difficult to exclude alternative hypotheses such as multiple gene losses ^30^. Here, should contamination underly the presence of plant-like genes in *B. tabaci*, it would have had to occur at least three times independently during the sequencing process of three genomes (MEAM1, MED, SAS), and it would have had to involve three times the same plant genes, and these genes only. Regarding a possible bias in terms of taxonomic representation of the target protein database, we would like to emphasize that the UniprotKB database, from which the Uniref90 database derives, is now particularly dense and diverse in terms of animal proteins. By browsing the “Proteomes” page of the “UniProt” website (https://www.uniprot.org/proteomes/), we found that it currently contains 1,093 animal proteomes comprising more than 10,000 proteins that are distributed among seven large clades (Chordata, Cnidaria, Ecdysozoa, Echinodermata, Placozoa, Porifera, Spiralia). Among Ecdysozoa, the database contains 151 insect proteomes that comprise at least 10,000 proteins and are distributed in eight orders (Coleoptera, Dictyoptera, Diptera, Hemiptera, Hymenoptera, Lepidoptera, Psocodea, Thysanoptera). It is thus unlikely that the narrow taxonomic distribution of *B. tabaci* plant-like genes we observe in animal proteomes (i.e. specific to Aleyrodidae) is due to a bias in the representativeness of animals or insect taxa in the target database. Furthermore, the gene loss alternative hypothesis would posit that *B. tabaci* plant-like genes were inherited vertically from a common ancestor with plants (i.e. the last eukaryote common ancestor ^31^) and were lost in all eukaryotes but plants and B. tabaci which, we believe, is highly unlikely. Regarding the possibility of phylogenetic artefact, we acknowledge that the branching of plant-like *B. tabaci* genes in our phylogenies may not be always strongly supported. However, given the absence of these genes in non-Aleyrodidae animals and their close phylogenetic proximity to plant genes, their acquisition through HGT seems more parsimonious to envision from a plant than a non-plant source.

### Inference of HGT and gene duplication events in Aleyrodidae

We next examined the extent to which these plant gene transfers have contributed to the genome of other Aleyrodidae available in Genbank. We clustered the 24 representative MEAM1 proteins with the whole proteomes from the *B. tabaci* cryptic species MEAM1, SSA-ECA and MED, and from *Trialeurodes vaporariorum*, the only other Aleyrodidae available in Genbank at the time we performed this study. In total, the 24 corresponding Aleyrodidae clusters encompass 138 proteins, of which 131 have best hits against plant homologs including 45 from SSA-ECA, 35 from MED, and 49 from MEAM1 *B. tabaci* cryptic species as well as two from *T. vaporarium* (Table 1, Supplementary Table 2). Protein clustering combined to phylogenetic analysis shows that at least 24 independent HGT events are necessary to explain the presence of these genes in *B. tabaci* genomes. This number of plant-to-insect HGT is remarkable given that most HGT events reported so far in animals (excluding HT of transposable elements) involve genes of bacterial or fungal origins ^32 18^, with only few cases of gene transfers from non-fungal eukaryotes to animals ^33 34^. It is noteworthy that fifteen out of twenty clusters are shared between the MEAM1 and the SSA-ECA cryptic species, indicating that most HGTs likely occurred before the split of these species, i.e., between 19 and 40 million years ago ^35,36.^

Several transfers were apparently followed by gene duplications. The largest cluster, with predicted delta(12) fatty acid desaturase (FAD) function, comprises 38 Aleyrodidae members including 15 MEAM1 proteins. Eight of the FAD genes are organized in a genomic region spanning about 120kb (scaffold 995: 912024-1039680) which is also observed in syntenic position at least in the SSA-ECA genome (scaffold 436:909978-1039696), indicating that this hotspot evolved before the split of these species. In combination with phylogenetic analysis, the amplification of the FAD genes can be explained by a mixture of local and distal duplication events post HGT (Figure 1A).

We notice, however, that in the FAD tree, the plant-related *B. tabaci* proteins are distributed on two distinct branches. In addition, the FAD tree is one of two clusters comprising a *T. vaporarium* protein, which appears more closely related to plant than to *B. tabaci* proteins (Figure 2A). We reasoned that this tree topology could be biased by the hit sampling procedure which uses only the MEAM1 cluster representatives as queries. To address the monophyly of the *Aleyrodidae* proteins, we therefore incorporated to the initial protein alignment a collection of additional Uniref proteins collected as for the MEAM1 representative proteins above. These proteins were retrieved from similarity searches using the *T. vaporarium* protein (Tv08972) and a representative of the second *B. tabaci* branch (Bta04264) as queries against the Uniref databases used in this study (see above and methods). This incorporation of additional hits from the target database is meant to refine the position of Tv08972 and Bta04264 in the tree. Remarkably, the resulting tree indicates monophyly of the *B. tabaci* proteins and confirms that the *T. vaporarium* protein is more closely related to plant proteins than to *B. tabaci* ones (Figure 3A). This suggests that independent HGT events of plant FAD genes occurred in *T. vaporarium* and in the common ancestor of the three *B. tabaci* cryptic species.

**Figure 3.**
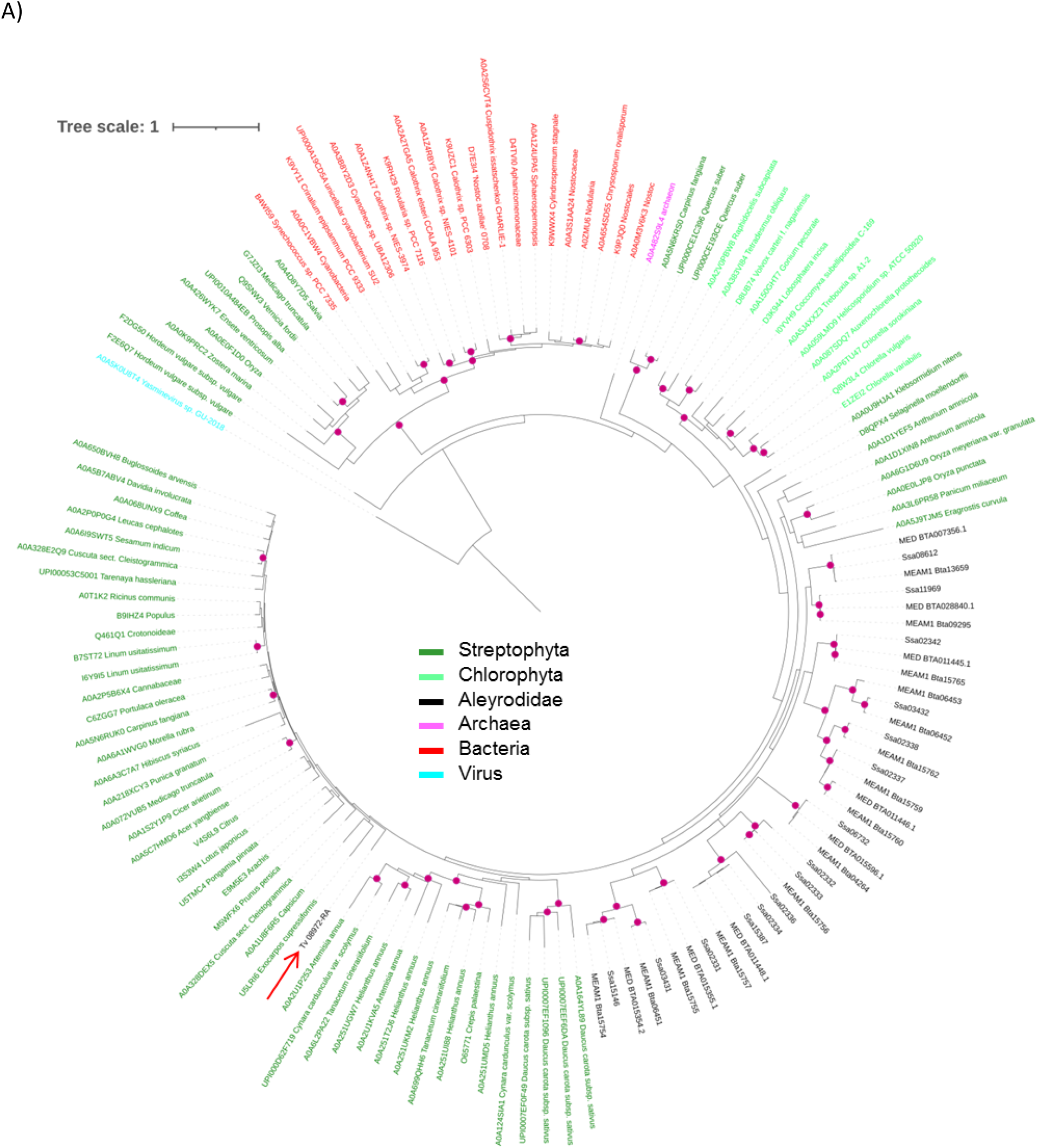
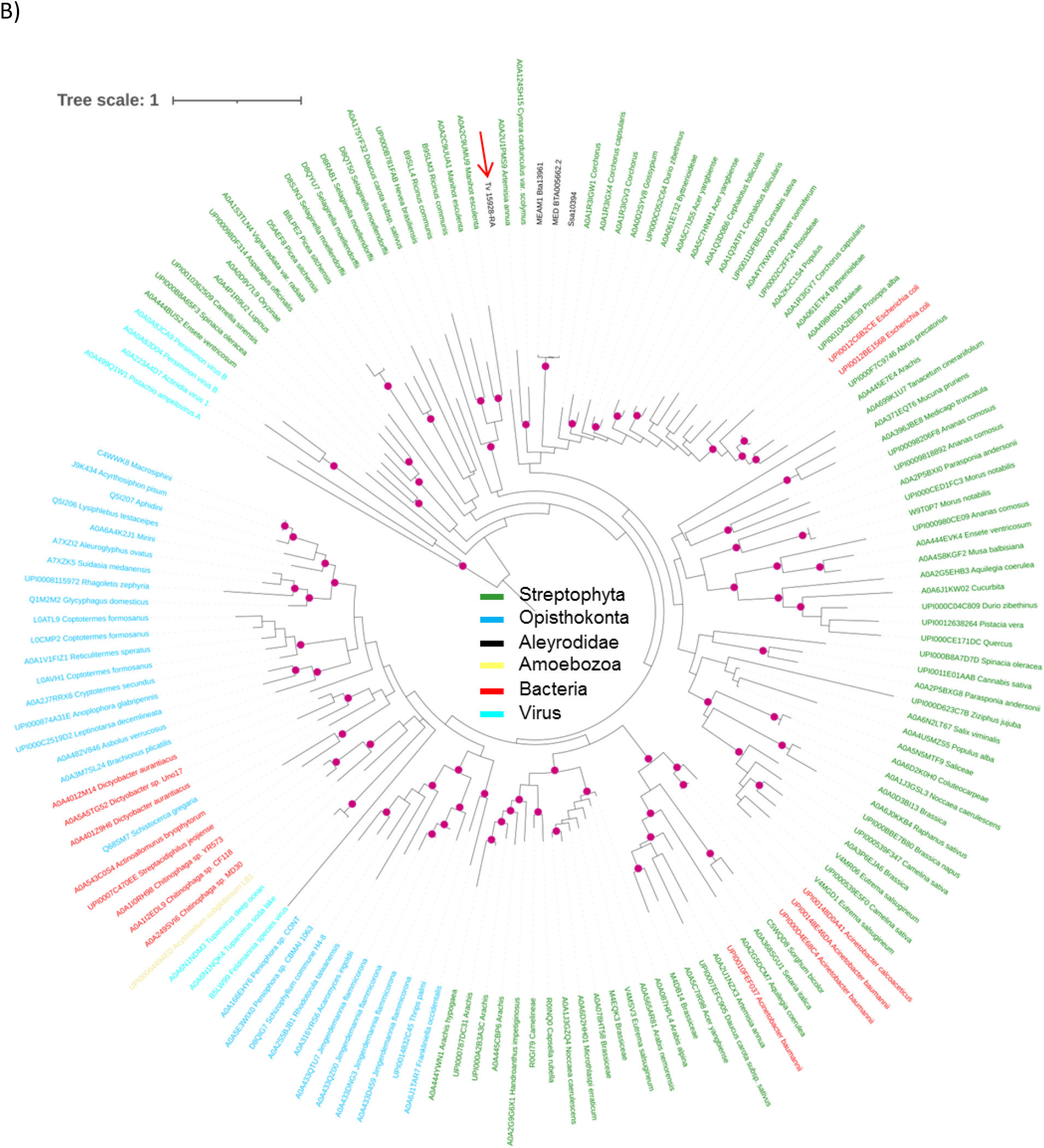
Phylogenetic relationships of potential plant-derived genes in Aleyrodidae. Phylogenetic trees showing the Bta06453 cluster, annotated as delta(12) fatty acid desaturase family (A) and the Bta13961 cluster, annotated as Thaumatin (B). In (A), the tree is inferred from the multiple sequence alignment of the cluster representative sequence (MEAM1 Bta06453), MEAM1_Bta04264 and Tv_08972 with their homologs found across Aleyrodidae proteins and Uniref90 proteins from eukaryotes, prokaryotes and viruses. In (B) the tree is inferred from the multiple sequence alignment of the cluster representative sequence (MEAM1 Bta13961) and Tv_15928 with their homologs found across Aleyrodidae proteins and Uniref90 proteins from eukaryotes, prokaryotes and viruses. The labels for Uniref90 proteins are either species names or the names of the first taxonomic level comprising all the proteins of a specific Uniref90 cluster. The SSA-ECA proteins are labelled as " Ssa ». The *Trialeurodes vaporariorum* proteins are labelled as “Tv” and indicated by a red arrow. The bootstrap values above 70% are indicated as magenta disks. The color legend indicating the taxonomy of proteins is indicated inset. For the ease of representation, the trees are presented in a circular mode rooted at midpoint. Unrooted trees are available in the Supplementary Figures 1 & 2.

From the initial phylogenetic tree of the second cluster including a *T. vaporarium* protein (Tv15928 in cluster Bta13961), the monophyly between the latter and the *B. tabaci* orthologs is also not supported. We performed the same data collection as for the FAD tree to enrich the Bta13961 protein alignment with hits of the *T. vaporarium* protein against the target databases. We observe that, again, phylogenetic clustering does not support monophyly at the level of Aleyrodidae, suggesting that an independent HGT event occurred in *T. vaporarium* (Figure 3B). Here however, the *T. vaporarium* protein does not group with close plant orthologs. Further supporting possible independent HGT in *T. vaporarium* and *B. tabaci* for these genes, we compared flanking protein coding genes between *T. vaporarium* and *B. tabaci* MEAM1 and were unable to detect any synteny between the two species (data not shown).

### Number of introns and codon usage bias of horizontally transferred genes

We addressed whether the structure of plant-derived genes could inform us regarding the nature of the transferred material. Indeed, genes are transferred with their introns when DNA is captured from the donor genome while they are not in the case of the capture of mRNA-derived retrogenes ^37^. Based on the gff3 annotation files of the three *B. tabaci* genomes, we found that plant-like genes in these genomes have an average of 1.8 introns, with 28 genes having zero predicted intron and 2 genes having a maximum of six predicted introns (Supplementary Table 2). Therefore, we could not observe strong evidence for RNA-based transfer and the presence of introns in most plant-derived genes could be interpreted as reflecting a DNA-based transfer or an RNA-mediated mechanism followed by intron acquisitions in Aleyrodidae genomes.

We also used the COUSIN index to assess whether *B. tabaci* plant-like genes showed any particular bias in codon usage that may further inform us on the way they have evolved after transfer ^38^ (https://cousin.ird.fr/). Given the age of the transfers and the relatively poorly resolved phylogenetic relationships of *B. tabaci* plant-like genes among plant genes, it was not possible to identify donor plant species. Thus, we could not compare the codon preference usage (CUPref) between donor and receiving (*B. tabaci*) species as in ^39^ for example. We plotted the distribution of COUSIN indexes of plant-like *B. tabaci* genes and compared it to that of all *B. tabaci* genes. We found that a substantial fraction of the *B. tabaci* CDS have a codon usage preference (CUPref) that differs from the null hypothesis, i.e. the distribution of COUSIN indexes is skewed toward scores of 1 or above (Supplementary Figure 2). However, most *B. tabaci* plant-like genes fall among genes that have no CUPref, i.e., the distribution of their COUSIN index is centered on zero (Supplementary Figure 2). One could propose that absence of CUPref may have facilitated incorporation of these genes in the cellular environment of *B. tabaci*. However, it is unclear whether these genes have no CUPref in the donor species or whether they evolved towards absence of CUPref after being transferred in the *B. tabaci* genome.

### Putative function of horizontally transferred genes

To search for evidence of functionality of plant-like genes in *B. tabaci*, we measured the evolutionary pressures acting on them by calculating pairwise ratios of non-synonymous over synonymous substitutions (Ka/Ks) between sequences within each of the 24 clusters. Importantly, these pairwise Ka/Ks ratio are here calculated within each cluster both between paralogs and between orthologs (between *B. tabaci* species). We found that 593 out of the 612 ratios we obtained were lower than 0.5, indicating that most if not all plant-like genes have evolved under purifying selection in the *B. tabaci* species complex. Furthermore, we found transcripts supporting expression of at least one gene for 18 out of the 24 clusters, often in multiple independent transcriptome assemblies (Table 1). Together with conservation across cryptic species and evidence of gene duplication, these data suggest that most if not all *B. tabaci* plant-like genes are functional.

Interestingly, most of these genes (21 out of 24 clusters) have a predicted function based on similarity with their nearest plant relative. As for the recently described malonyltransferase ^24,40^, there is direct or indirect evidence that many of them are involved in plant-pathogen interactions (Table 1). For example, the delta(12) FAD are known to produce polyunsaturated linoleic acid in plants, which are involved in response to pathogens ^41^. Likewise, members of subtilisin-like protease and pathogen-related protein families can both be induced following pathogen infection ^42,43^. Ornithine decarboxylase is also worth noting as this enzyme synthesizes putrescine in plants, a polyamine involved in pathogen response ^44^ and suspected to be usurped by Hessian fly larvae to facilitate nutrient acquisition while feeding on wheat ^45^. In the same vein, a gene resembling the nicotianamine synthase, involved in the transport of various metal ions in plants ^46^, may facilitate acquisition of micronutrient by whiteflies. Finally, pectinesterase is a plant cell wall degrading enzyme (PCWDE) that is also found in plant and fungal pathogens causing maceration and soft-rotting of plant tissues. In fact, horizontal acquisition of PCWDE has already been documented in insects, but the source of the gene was bacterial ^47^.

To conclude, our study reveals that in addition to bacterial genes, which repeatedly entered arthropod genomes and fueled the evolution of herbivory ^18,48^, numerous plant genes have been acquired through HGT by *B. tabaci*, likely contributing to the highly polyphagous nature of this species. The significant representation of predicted functions potentially involved in parasitism suggests that these genes were selected from an important set of transferred genes. Using the same approach on *Drosophila melanogaster*, we found no gene of potential plant origin, showing that plant-to-insect HGT is not ubiquitous and suggesting that it could be facilitated by specific vectors in association with *B. tabaci*. It is noteworthy that viruses have been proposed to act as vectors of HT in eukaryotes ^49,50^ and that *B. tabaci* is known to act as a vector of dozens of plant viruses, some of which are able to replicate in insect cells ^51^. Our results call for a large-scale evaluation of plant-to-insect HGT and for a detailed functional characterization of *B. tabaci* plant-like genes, which may further contribute to control this pest.

## Materials and Methods

### Protein and genome databases

The genomes, proteomes and CDS from *T. vaporariorum* and the *Bemisia tabaci* cryptic species MEAM1, SSA-ECA and MED were retrieved from the whitefly database (http://www.whiteflygenomics.org/ftp/). The Uniref90 (U90) database was downloaded on February 1^st^ 2021 (https://www.uniprot.org/downloads).

### Prediction of plant-derived genes in Aleyrodidae

We used MMseqs2 taxonomy ^26^ with three different modes of LCA inference (--lca-mode 1, 2 and 4) to infer taxonomic origin of *B. tabaci* MEAM1 predicted proteins against Uniref90 (U90) database from which Aleyrodidae proteins were excluded. The sequences inferred as of Viridiplantae origin were selected when ancestry in plants could be established deeper than genus level, i.e. at least at the family level. The 65 MEAM1 proteins meeting this criterion were clustered using MMseqs when identity between query and target was above 35% and when target coverage by query was at least 30% (-c 0.3 --cov-mode 1 --min-seq-id 0.35) resulting in 41 clusters from which representative sequences were selected. The predicted proteins from *T. vaporariorum*, and B. tabaci MEAM1, SSA-ECA and MED cryptic species were clustered with the same parameters and the sequences from clusters containing MEAM1 representative plant proteins were selected. Each of these Aleyroridae proteins was compared to U90 to confirm or not their plant origin on the basis of best hit taxonomy (i.e. Viridiplantae or other).

### Construction of phylogenetic trees

Representative MEAM1 sequences were used to search for homologs with MMseqs2 search module against the following five databases. Four are taxonomic subsets of the U90 database corresponding to proteins from (1) Streptophyta; (2) eukaryotes without Steptophyta and without Aleyrodidae; (3) prokaryotes; (4) viruses. The fifth database comprises Aleyrodidae proteomes from *B. tabaci* cryptic species MEAM1, MED and SSA-ECA and from *Trialeurodes vaporariorum*. For each query, the fasta sequences of the top ten hits obtained against each database supplemented with the twenty (40 in the case of Bta13961) most diverse hits obtained against Streptophyta were retrieved and aligned using MAFFT v7.475 (local alignment) ^52^. Poorly aligned sequences were removed using trimal v1.4 (-resoverlap 0.7 -seqoverlap 70) ^53^ unless this removes the MEAM1 proteins in which case all sequences were kept. The selected sequences were aligned again with MAFFT (local alignment) and the less informative columns of the alignments were removed using trimal (-gappyout). The resulting alignments were used for model selection and phylogenetic reconstruction using IQTREE v1.6.12 ^54^ with 100 bootstrap replicates.

### Evidence of functionality

To assess under which selective regime plant-like genes have evolved after transfer in *B. tabaci*, we aligned *B. tabaci* sequences within each cluster at the codon level using MACSE v2 ^55^, trimmed the codon alignment using trimal v1.4 (-backtrans -ignorestopcodon -gt 0.8) ^53^ and calculated ratios of non-synonymous substitutions over synonymous substitution Ka/Ks between all pairs of sequences using the seqinr R package ^56^. The number of ratios below 0.5 (indicating purifying selection) was counted and is reported in Table 1 and Supplementary Table 2. We also searched for evidence that *B. tabaci* plant-like genes are transcribed by using all sequences from the MEAM1 cryptic species as queries to perform blastn (-task megablast) searches on 12 *B. tabaci* MEAM transcriptomes retrieved from Genbank under the following accession numbers: GAPP01.1, GAPQ01.1, GARP01.1, GARQ01.1, GAUC01.1, GBII01.1, GBIJ01.1, GCZW01.1, GEZK01.1, GFXM01.1, GIBX01.1, GICC01.1. All transcripts aligning over at least 75% of the length of the *B. tabaci* coding sequence with at least 95% nucleotide identity were counted and reported in Supplementary Table 2. Accession number and alignment features of these transcripts are provided in Supplementary Table 3.

## Data availability

Supplementary Dataset (on [figshare repository](https://figshare.com/projects/Gilbert_and_Maumus_-_Supplementary_Dataset/128792)) containing

- Supplementary table 1 providing the best hits of the proteins coded by the genes flanking the plant-derived genes in MEAM1.
- Supplementary table 2 providing features of all *B. tabaci* plant-derived genes included in this study and references supporting involvement in plant-pathogen interactions.
- Supplementary table 3 listing all transcripts covering at least 75% of the *B. tabaci* MEAM1 plant-derived genes with at least 95% identity.
- Supplementary Dataset 1, the LCA inference reports for the three taxonomy modes
- Supplementary Dataset 2, an archive an archive comprising the initial (.aln files) and trimmed (.trim files) protein alignments in fasta format as well as the phylogenetic trees in Newick format (.annot files) for each cluster of false positives.
- Supplementary Dataset 3, an archive comprising the initial (.aln files) and trimmed (.trim files) protein alignments in fasta format as well as the phylogenetic trees in Newick format (.annot files) for each cluster of plant-derived genes.
- Supplementary Dataset 4, a fasta file combining the plant-derived MEAM1 representative sequences
- Supplementary Figure 1, the tree images combined in a single file for the false positives (.pdf)
- Supplementary Figure 2, the tree images combined in a single file for plant-derived genes (.pdf).
- Supplementary Figure 3, the figure presenting the result of the COUSIN analysis.

All the phylogenetic trees can be viewed interactively at https://itol.embl.de/shared/fmaumus

## Notes

**Competing interest statement:** All authors declare to have no conflict of interest

https://figshare.com/projects/Gilbert_and_Maumus_-_Supplementary_Dataset/128792

### Competing Interest Statement

The authors have declared no competing interest.

## References

1 Soucy, S. M., Huang, J. & Gogarten, J. P. Horizontal gene transfer: building the web of life. Nat Rev Genet 16, 472–482, doi:10.1038/nrg3962 (2015).

2 Martin, W. F. Too Much Eukaryote LGT. Bioessays 39, doi:10.1002/bies.201700115 (2017).

3 Salzberg, S. L. Horizontal gene transfer is not a hallmark of the human genome. Genome Biol 18, 85, doi:10.1186/s13059-017-1214-2 (2017).

4 Van Etten, J. & Bhattacharya, D. Horizontal Gene Transfer in Eukaryotes: Not if, but How Much? Trends Genet 36, 915–925, doi:10.1016/j.tig.2020.08.006 (2020).

5 Sibbald, S. J., Eme, L., Archibald, J. M. & Roger, A. J. Lateral Gene Transfer Mechanisms and Pan-genomes in Eukaryotes. Trends Parasitol 36, 927–941, doi:10.1016/j.pt.2020.07.014 (2020).

6 Husnik, F. & McCutcheon, J. P. Functional horizontal gene transfer from bacteria to eukaryotes. Nat Rev Microbiol 16, 67–79, doi:10.1038/nrmicro.2017.137 (2018).

7 Leger, M. M., Eme, L., Stairs, C. W. & Roger, A. J. Demystifying Eukaryote Lateral Gene Transfer (Response to Martin 2017 DOI: 10.1002/bies.201700115). Bioessays 40, e1700242, doi:10.1002/bies.201700242 (2018).

8 Lacroix, B. & Citovsky, V. Transfer of DNA from Bacteria to Eukaryotes. mBio 7, doi:10.1128/mBio.00863-16 (2016).

9 Hibdige, S. G. S., Raimondeau, P., Christin, P. A. & Dunning, L. T. Widespread lateral gene transfer among grasses. New Phytol 230, 2474–2486, doi:10.1111/nph.17328 (2021).

10 Yang, Z. et al. Convergent horizontal gene transfer and cross-talk of mobile nucleic acids in parasitic plants. Nat Plants 5, 991–1001, doi:10.1038/s41477-019-0458-0 (2019).

11 Cai, L. et al. Deeply Altered Genome Architecture in the Endoparasitic Flowering Plant Sapria himalayana Griff. (Rafflesiaceae). Curr Biol 31, 1002–1011 e1009, doi:10.1016/j.cub.2020.12.045 (2021).

12 Paganini, J. et al. Contribution of lateral gene transfers to the genome composition and parasitic ability of root-knot nematodes. PLoS One 7, e50875, doi:10.1371/journal.pone.0050875 (2012).

13 Simion, P. et al. Chromosome-level genome assembly reveals homologous chromosomes and recombination in asexual rotifer Adineta vaga. Sci Adv 7, eabg4216, doi:10.1126/sciadv.abg4216 (2021).

14 Dunning Hotopp, J. C. Horizontal gene transfer between bacteria and animals. Trends Genet 27, 157–163, doi:10.1016/j.tig.2011.01.005 (2011).

15 Cummings, T. F. M. et al. Citrullination Was Introduced into Animals by Horizontal Gene Transfer from Cyanobacteria. Mol Biol Evol 39, doi:10.1093/molbev/msab317 (2022).

16 Verster, K. I. et al. Horizontal Transfer of Bacterial Cytolethal Distending Toxin B Genes to Insects. Mol Biol Evol 36, 2105–2110, doi:10.1093/molbev/msz146 (2019).

17 Moran, N. A. & Jarvik, T. Lateral transfer of genes from fungi underlies carotenoid production in aphids. Science 328, 624–627, doi:10.1126/science.1187113 (2010).

18 Wybouw, N., Pauchet, Y., Heckel, D. G. & Van Leeuwen, T. Horizontal Gene Transfer Contributes to the Evolution of Arthropod Herbivory. Genome Biol Evol 8, 1785–1801, doi:10.1093/gbe/evw119 (2016).

19 Haegeman, A., Jones, J. T. & Danchin, E. G. Horizontal gene transfer in nematodes: a catalyst for plant parasitism? Mol Plant Microbe Interact 24, 879–887, doi:10.1094/MPMI-03-11-0055 (2011).

20 Danchin, E. G., Guzeeva, E. A., Mantelin, S., Berepiki, A. & Jones, J. T. Horizontal Gene Transfer from Bacteria Has Enabled the Plant-Parasitic Nematode Globodera pallida to Feed on Host-Derived Sucrose. Mol Biol Evol 33, 1571–1579, doi:10.1093/molbev/msw041 (2016).

21 Acuna, R. et al. Adaptive horizontal transfer of a bacterial gene to an invasive insect pest of coffee. Proc Natl Acad Sci U S A 109, 4197–4202, doi:10.1073/pnas.1121190109 (2012).

22 Wybouw, N. et al. A gene horizontally transferred from bacteria protects arthropods from host plant cyanide poisoning. Elife 3, e02365, doi:10.7554/eLife.02365 (2014).

23 Lapadula, W. J., Mascotti, M. L. & Juri Ayub, M. Whitefly genomes contain ribotoxin coding genes acquired from plants. Sci Rep 10, 15503, doi:10.1038/s41598-020-72267-1 (2020).

24 Xia, J. et al. Whitefly hijacks a plant detoxification gene that neutralizes plant toxins. Cell 184, 1693–1705 e1617, doi:10.1016/j.cell.2021.02.014 (2021).

25 Chen, W. et al. The draft genome of whitefly Bemisia tabaci MEAM1, a global crop pest, provides novel insights into virus transmission, host adaptation, and insecticide resistance. BMC Biol 14, 110, doi:10.1186/s12915-016-0321-y (2016).

26 Mirdita, M., Steinegger, M., Breitwieser, F., Soding, J. & Levy Karin, E. Fast and sensitive taxonomic assignment to metagenomic contigs. Bioinformatics, doi:10.1093/bioinformatics/btab184 (2021).

27 Suzek, B. E. et al. UniRef clusters: a comprehensive and scalable alternative for improving sequence similarity searches. Bioinformatics 31, 926–932, doi:10.1093/bioinformatics/btu739 (2015).

28 UniProt, C. UniProt: a worldwide hub of protein knowledge. Nucleic Acids Res 47, D506–D515, doi:10.1093/nar/gky1049 (2019).

29 Bemm, F., Weiss, C. L., Schultz, J. & Forster, F. Genome of a tardigrade: Horizontal gene transfer or bacterial contamination? Proc Natl Acad Sci U S A 113, E3054–3056, doi:10.1073/pnas.1525116113 (2016).

30 Dunning Hotopp, J. C. Grafting or pruning in the animal tree: lateral gene transfer and gene loss? BMC Genomics 19, 470, doi:10.1186/s12864-018-4832-5 (2018).

31 Burki, F., Roger, A. J., Brown, M. W. & Simpson, A. G. B. The New Tree of Eukaryotes. Trends Ecol Evol 35, 43–55, doi:10.1016/j.tree.2019.08.008 (2020).

32 Wybouw, N., Van Leeuwen, T. & Dermauw, W. A massive incorporation of microbial genes into the genome of Tetranychus urticae, a polyphagous arthropod herbivore. Insect Mol Biol 27, 333–351, doi:10.1111/imb.12374 (2018).

33 Graham, L. A. & Davies, P. L. Horizontal Gene Transfer in Vertebrates: A Fishy Tale. Trends Genet 37, 501–503, doi:10.1016/j.tig.2021.02.006 (2021).

34 Gladyshev, E. A., Meselson, M. & Arkhipova, I. R. Massive horizontal gene transfer in bdelloid rotifers. Science 320, 1210–1213, doi:10.1126/science.1156407 (2008).

35 Santos-Garcia, D., Vargas-Chavez, C., Moya, A., Latorre, A. & Silva, F. J. Genome evolution in the primary endosymbiont of whiteflies sheds light on their divergence. Genome Biol Evol 7, 873–888, doi:10.1093/gbe/evv038 (2015).

36 Mugerwa, H. et al. African ancestry of New World, Bemisia tabaci-whitefly species. Sci Rep 8, 2734, doi:10.1038/s41598-018-20956-3 (2018).

37 Brosius, J. Retroposons--seeds of evolution. Science 251, 753, doi:10.1126/science.1990437 (1991).

38 Bourret, J., Alizon, S. & Bravo, I. G. COUSIN (COdon Usage Similarity INdex): A Normalized Measure of Codon Usage Preferences. Genome Biol Evol 11, 3523–3528, doi:10.1093/gbe/evz262 (2019).

39 Callens, M., Scornavacca, C. & Bedhomme, S. Evolutionary responses to codon usage of horizontally transferred genes in Pseudomonas aeruginosa: gene retention, amelioration and compensatory evolution. Microb Genom 7, doi:10.1099/mgen.0.000587 (2021).

40 Taguchi, G. et al. Malonylation is a key reaction in the metabolism of xenobiotic phenolic glucosides in Arabidopsis and tobacco. Plant J 63, 1031–1041, doi:10.1111/j.1365-313X.2010.04298.x (2010).

41 Dar, A. A., Choudhury, A. R., Kancharla, P. K. & Arumugam, N. The FAD2 Gene in Plants: Occurrence, Regulation, and Role. Front Plant Sci 8, 1789, doi:10.3389/fpls.2017.01789 (2017).

42 van Loon, L. C., Rep, M. & Pieterse, C. M. Significance of inducible defense-related proteins in infected plants. Annu Rev Phytopathol 44, 135–162, doi:10.1146/annurev.phyto.44.070505.143425 (2006).

43 Figueiredo, A., Monteiro, F. & Sebastiana, M. Subtilisin-like proteases in plant-pathogen recognition and immune priming: a perspective. Front Plant Sci 5, 739, doi:10.3389/fpls.2014.00739 (2014).

44 Rhee, H. J., Kim, E. J. & Lee, J. K. Physiological polyamines: simple primordial stress molecules. J Cell Mol Med 11, 685–703, doi:10.1111/j.1582-4934.2007.00077.x (2007).

45 Subramanyam, S. et al. Hessian fly larval feeding triggers enhanced polyamine levels in susceptible but not resistant wheat. BMC Plant Biol 15, 3, doi:10.1186/s12870-014-0396-y (2015).

46 Laffont, C. & Arnoux, P. The ancient roots of nicotianamine: diversity, role, regulation and evolution of nicotianamine-like metallophores. Metallomics 12, 1480–1493, doi:10.1039/d0mt00150c (2020).

47 Shelomi, M. et al. Horizontal Gene Transfer of Pectinases from Bacteria Preceded the Diversification of Stick and Leaf Insects. Sci Rep 6, 26388, doi:10.1038/srep26388 (2016).

48 Luan, J. B. et al. Metabolic Coevolution in the Bacterial Symbiosis of Whiteflies and Related Plant Sap-Feeding Insects. Genome Biol Evol 7, 2635–2647, doi:10.1093/gbe/evv170 (2015).

49 Gilbert, C. & Cordaux, R. Viruses as vectors of horizontal transfer of genetic material in eukaryotes. Curr Opin Virol 25, 16–22, doi:10.1016/j.coviro.2017.06.005 (2017).

50 Catoni, M. et al. Virus-mediated export of chromosomal DNA in plants. Nat Commun 9, 5308, doi:10.1038/s41467-018-07775-w (2018).

51 He, Y. Z. et al. A plant DNA virus replicates in the salivary glands of its insect vector via recruitment of host DNA synthesis machinery. Proc Natl Acad Sci U S A 117, 16928–16937, doi:10.1073/pnas.1820132117 (2020).

52 Katoh, K., Misawa, K., Kuma, K. & Miyata, T. MAFFT: a novel method for rapid multiple sequence alignment based on fast Fourier transform. Nucleic Acids Res 30, 3059–3066 (2002).

53 Capella-Gutierrez, S., Silla-Martinez, J. M. & Gabaldon, T. trimAl: a tool for automated alignment trimming in large-scale phylogenetic analyses. Bioinformatics 25, 1972–1973, doi:10.1093/bioinformatics/btp348 (2009).

54 Minh, B. Q. et al. IQ-TREE 2: New Models and Efficient Methods for Phylogenetic Inference in the Genomic Era. Mol Biol Evol 37, 1530–1534, doi:10.1093/molbev/msaa015 (2020).

55 Ranwez, V., Douzery, E. J. P., Cambon, C., Chantret, N. & Delsuc, F. MACSE v2: Toolkit for the Alignment of Coding Sequences Accounting for Frameshifts and Stop Codons. Mol Biol Evol 35, 2582–2584, doi:10.1093/molbev/msy159 (2018).

56 Charif, D. & Lobry, J. R. In Structural Approaches to Sequence Evolution: Molecules, Networks, Populations (eds Ugo Bastolla, Markus Porto, H. Eduardo Roman, & Michele Vendruscolo) 207–232 (Springer Berlin Heidelberg, 2007).

